# Genomic and transcriptomic somatic alterations of hepatocellular carcinoma in non-cirrhotic livers

**DOI:** 10.1101/2021.12.14.472689

**Authors:** Zachary L Skidmore, Jason Kunisaki, Yiing Lin, Kelsy C Cotto, Erica K Barnell, Jasreet Hundal, Kilannin Krysiak, Vincent Magrini, Lee Trani, Jason R Walker, Robert Fulton, Elizabeth M Brunt, Christopher A Miller, Richard K Wilson, Elaine R Mardis, Malachi Griffith, William Chapman, Obi L Griffith

## Abstract

**Background:** Liver cancer is the second leading cause of cancer-related deaths worldwide. Hepatocellular carcinoma (HCC) risk factors include chronic hepatitis, cirrhosis, and alcohol abuse, whereby tumorigenesis is induced through inflammation and subsequent fibrotic response. However, a subset of HCC arises in non-cirrhotic livers. We characterized the genomic and transcriptomic landscape of non-cirrhotic HCC to identify features underlying the disease’s development and progression.

**Methods:** Whole genome and transcriptome sequencing was performed on 30 surgically resectable tumors comprised of primarily of non-cirrhotic HCC and adjacent normal tissue. Using somatic variants, capture reagents were created and employed on an additional 87 cases of mixed cirrhotic/non-cirrhotic HCC. Cases were analyzed to identify viral integrations, single nucleotide variants (SNVs), insertions and deletions (INDELS), copy number variants, loss of heterozygosity, gene fusions, structural variants, and differential gene expression.

**Results:** We detected 3,750 SNVs/INDELS and extensive CNVs and expression changes. Recurrent *TERT* promoter mutations occurred in >52% of non-cirrhotic discovery samples. Frequently mutated genes included *TP53*, *CTNNB1*, and *APOB*. Cytochrome P450 mediated metabolism was significantly downregulated. Structural variants were observed at *MACROD2, WDPCP* and *NCKAP5* in >20% of samples. Furthermore, *NR1H4* fusions involving gene partners *EWSR1, GNPTAB*, and *FNIP1* were detected and validated in 2 non-cirrhotic samples.

**Conclusion:** Genomic analysis can elucidate mechanisms that may contribute to non-cirrhotic HCC tumorigenesis. The comparable mutational landscape between cirrhotic and non-cirrhotic HCC supports previous work suggesting a convergence at the genomic level during disease progression. It is therefore possible genomic-based treatments can be applied to both HCC subtypes with progressed disease.

**Highlights:**

- Non-cirrhotic HCC genomically resembles cirrhotic HCC
- Comprehensive genome- and transcriptome-wide profiling allows detection of novel structural variants, fusions, and undiagnosed viral infections
- *NR1H4* fusions may represent a novel mechanism for tumorigenesis in HCC
- Non-cirrhotic HCC is characterized by genotoxic mutational signatures and dysregulated liver metabolism
- Clinical history and comprehensive omic profiling incompletely explain underlying etiologies for non-cirrhotic HCC highlighting the need for further research

**Short Description:** This study characterizes the genomic landscape of hepatocellular carcinomas (HCCs) in non-cirrhotic livers. Using 117 HCCs tumor/normal pairs, we identified 3,750 SNVs/INDELS with high variant frequency in TERT, *TP53*, *CTNNB1*, and *APOB*. *CYP450* was significantly downregulated and many structural variants were observed. This characterization could assist in elucidating non-cirrhotic HCC tumorigenesis.

## Introduction

Worldwide, there are approximately 750,000 new cases of hepatocellular carcinoma (HCC) each year [1]. Although HCC has the 5th highest incidence rate in men and 9th highest incidence rate in women, it has the second highest mortality rate of all cancer types [1]. HCC is traditionally associated with inflammation-inducing risk factors, which promote liver cirrhosis including: chronic hepatitis infections, such as hepatitis b virus (HBV) and hepatitis c virus (HCV), alcohol abuse, and non-alcoholic fatty liver disease [2]. However, approximately 20% of patients present with non-cirrhotic HCC in the absence of these risk factors [3]. If diagnosed early, patients with non-cirrhotic HCC maintain adequate liver function, allowing for effective tumor resection with exceptional prognosis when compared to patients with cirrhotic HCC [4]. However, late-stage diagnosis of non-cirrhotic HCC typically presents with larger and more aggressive tumors that are prone to metastasis [5]. Even with extensive tumor resection, approximately 50% of patients relapse within three years post-treatment [6].

Using high-throughput sequencing, researchers have previously characterized the genomic landscape of cirrhotic HCC [7–13]. These studies included whole genome, whole exome, and/or transcriptome sequencing with a focus on analyzing HCC induced by HBV, HCV, and/or cirrhosis. Prior studies, which have evaluated the genomics of cirrhotic and non-cirrhotic HCC, report that among the most significant and recurrent alterations are *TERT* mutations which typically occur at the promoter region [8,9,14]. Mutations within this region have been observed in a variety of cancer types beyond cirrhotic HCC, suggesting a common role of activating *TERT* promoter variants in oncogenesis and metastasis [15–17]. *TERT* expression in terminally differentiated cells promotes telomere maintenance and elongation [18]. Telomere maintenance is required for late stage cancer propagation with *TERT* misregulation being harnessed by human cancers to evade mitotic catastrophe and apoptosis [19]. Previous studies have recognized that increases in *TERT* expression could serve as a proxy for telomere maintenance; however, late-stage tumors exhibit shortened telomeres in comparison to their normal counterparts, due to high turnover rates [20,21]. Among studies specific to cirrhotic HCC, the putative mechanisms of *TERT* activation can be divided into three categories: 1) HBV integration events in the *TERT* promoter [8,22], 2) point mutations (C228T and C250T) in the promoter region mutually exclusive of HBV integration [9,23], and 3) structural variations of the *TERT* promoter region [8,14].

This study characterizes biomarkers and elucidates recurrent anomalies in non-cirrhotic HCC. We identified somatic variants in 117 tumor samples whereby 52 samples were cirrhotic, 63 samples were non-cirrhotic, and 2 samples had an unspecified cirrhotic status. Using this cohort, we analyzed single nucleotide variants (SNVs), insertions and deletions (INDELs), structural variation (SV), copy number variation (CNV), loss of heterozygosity (LOH), differential expression, and viral integration events. This comprehensive approach uncovered the genomic features implicated in non-cirrhotic HCC to improve its diagnosis, prognosis, and treatment.

## Methods

Refer to supplementary methods for more details.

### Sample Procurement

The discovery cohort consisted of 30 primary tumor and adjacent matched non-tumor liver samples obtained through surgical resection from adult patients diagnosed with HCC between 2000 to 2011 at the Washington University School of Medicine. Within this cohort, 13 were male and 17 were female. Additionally, 2 were African American and 28 were Caucasian. None of these samples exhibited evidence of hepatocellular adenoma (HCA) and the non-cirrhotic samples did not show signs of advanced fibrosis. 1 sample was HBV positive and 4 samples were HCV positive according to clinical data. All other samples within the discovery cohort had an unknown clinical etiology. The extension-alpha and extension-beta cohorts had 16 HCC tumors with matched non-tumor liver and 71 tumor-only HCC samples, respectively. Discovery and extension-alpha cohort samples were flash-frozen prior to banking and extension-beta samples were derived from formalin fixed paraffin embedded (FFPE) blocks. Across both extension cohorts, 27 were female and 58 were male. Furthermore, 2 were Asian, 13 were African American, and 70 were Caucasian. Within the extension-alpha cohort, two samples were HCV positive, one had chronic cholestasis, and the others had no known clinical etiology. Clinical data for the extension-beta cohort was as follows: 5 had known alcohol use, 8 were HBV positive, 29 were HCV positive, 2 were diagnosed with primary sclerosing cholangitis (PSC), and 6 samples were diagnosed with non-alcoholic steatohepatitis (NASH). From the extension-alpha cohort, 2 patients did not provide information on race and gender (**Table S1**). All patient samples were acquired after informed consent to an approved study by the Washington University School of Medicine Institutional Review Board (IRB 201106388).

### Sample Preparation and Sequencing

DNA and RNA from samples in the discovery cohort were extracted using the QIAamp DNA Mini kit and Qiagen RNeasy Mini kit, respectively. Whole genome sequencing libraries were constructed using Kapa HYPER kits for use on the Illumina HiSeq 2000 platform. The Ovation RNA-seq System V2 (NuGen Inc) kit was used to generate RNAseq libraries. Resulting barcoded libraries were pooled prior to Illumina sequencing. To validate variants identified from WGS, a hybrid capture panel (CAP1) was designed and executed on the Illumina platform to capture fragments from the WGS libraries. The QIAamp DNA Mini kit was used to extract DNA from extension-alpha samples, which was subsequently sequenced using the CAP1 strategy. Finally, CAP1 sequencing was used to identify variants from the DNA extracted from extension-beta samples with the QIAamp DNA FFPE Tissue kit. A second hybrid capture panel (CAP2) utilized Nimblegen and spiked-in IDT probes that hybridized to the *TERT* promoter locus and HBV genome (designed against a consensus sequence for 10 common HBV strains, see supplementary methods). CAP2 sequencing was employed on all 117 samples. *TERT* promoter variants were also detected in the discovery and extension-alpha cohorts with Sanger sequencing. cDNA capture was performed on pooled samples from the extension cohorts.

### Sequencing Alignment

WGS and CAP1 data were aligned to GRCh37 via the Genome Modeling System (GMS) using BWA [24,25]. Reads from the CAP2 data were competitively aligned using BWA [25] against the human reference genome (GRCh37) along with ten HBV genotypes for which complete genomes were available. RNAseq data were aligned with bowtie/tophat and expression was evaluated with cufflinks [26,27]. All raw RNAseq reads from the discovery cohort were also aligned against the HBV genomes for evidence of HBV expression at the RNA level. The predominant HBV strain was determined using relative coverage for competitive alignments. The precise location of the HBV integration site was identified from discordant read pairs from realigning HBV CAP2 reads to GRCh37 and the predominant HBV strain’s genome. A similar procedure was performed for HCV whereby both WGS and RNAseq reads were aligned against six HCV genotypes. The predominant HCV strain was determined using the total read support. To detect AAV1 and AAV2 integration, RNAseq reads were competitively aligned using kallisto [28] against AAV1 and AAV2 sequences.

### Telomere Length Determination

Telomeric tumor:normal read ratios were determined from WGS data using the GMS and visualized in R. A Wilcoxon-Mann-Whitney test measured the significance of differences between telomere length in tumor and normal samples.

### Variant Calling

Somatic variant analysis for single nucleotide variants (SNV) and insertions/deletions (INDEL) were performed on all three cohorts while germline variant analysis for these variants was performed on the discovery and extension-alpha cohort. Several computational tools within and outside of the GMS [29] were employed to facilitate variant calling and subsequent filtering based on variables including variant allele frequency, read count, and predicted pathogenicity.

### Structural Variant, Copy Number Variant, and Loss of Heterozygosity Analysis

WGS data from samples within the discovery cohort were analyzed for structural variants (SV), copy number variation (CNV), and loss of heterozygosity (LOH). Manta [30] was used to identify SV events. Manta-reported breakpoints, along with a 10kb flank were annotated with biomaRt and ensembl (GRCh37.p13). Regions of CNV were identified with the GMS and LOH were identified using VarScan2 [29,31]. The DNAcopy circular binary segmentation algorithm generated segments of LOH and CNV, which served as input for GISTIC [32] to conduct a recurrence analysis.

### NanoString nCounter Elements^tm^ Tagsets: *NR1H4* Fusion Validation

Fusion detection algorithms identified samples in the discovery cohort harboring gene fusions from RNAseq data. Fusion predictions involving *NR1H4* were validated across all 117 samples using a NanoString nCounter^®^ Elements™ TagSets assay. Sequences for predicted transcripts of the fusion calls that met certain read support criteria (≥10 spanning + encompassing reads and ≥1 spanning read) were sent to NanoString for probe design.

### Survival and Clinical Analysis

The R “survival” package [33] was used to associate SV-affected genes and CNV/LOH-affected genomic regions with overall survival and recurrence free survival. Only mutated genes and genomic regions occuring in ≥ 4 discovery cohort samples were included in this analysis. A survival analysis was also applied to SNV/INDELs observed in all non-cirrhotic samples from the three cohorts. All Kaplan-Meier survival plots were created in R. Fisher’s exact test was used to test for clinical associations with variables: lymphovascular space invasion (LVSI), tumor differentiation status, cirrhosis, and liver disease. Samples without relevant clinical data were excluded. Significance was measured with a multiple test correction using the FDR methodology (q-value < 0.05).

### Differential Expression and Pathway Analysis

Read counts for genes mutated in non-cirrhotic tumors and matched normal samples of the discovery cohort were used by the DEseq2 Bioconductor package [34] to perform differential gene expression analysis using a negative binomial distribution with samples as a blocking factor. Significance was measured with a Wald test and Benjamini & Hochberg multiple test correction (q-value < 0.5). Pathway analysis was performed using log2 differential expression data.

## Results

### Discovery Cohort

There were 30 patients included in the discovery cohort with tumors which were surgically resectable. These surgically resectable tumors were untreated, providing the opportunity to study HCC in the absence of chemotherapeutic intervention, which is normally incorporated in the treatment of cirrhotic HCC. Three of the patients within this cohort developed HCC in the setting of cirrhosis, all of which had been previously diagnosed with HCV. The remaining 27 individuals developed non-cirrhotic HCC, two of these individuals were diagnosed with HBV and another two individuals were diagnosed with HCV. To elucidate the genomic landscape of resected, primarily non-cirrhotic HCC, we performed whole genome sequencing (WGS), hybrid capture sequencing (CAP1), and transcriptome sequencing (RNAseq) on these 30 samples (**Table 1**). WGS failed for one tumor sample in the discovery cohort, therefore the final data for this cohort included WGS and CAP1 data for 29 samples (26 non-cirrhotic, 3 cirrhotic), and RNAseq data for 30 samples (27 non-cirrhotic, 3 cirrhotic). The sequencing analysis revealed a single previously unknown and undiagnosed HBV case with viral integration occurring at the *TERT* promoter (**Figure S1**, **Table S1**). Median haploid coverage for WGS data was 35.6x (range: 28.5-39.3) and 58.4x (range: 46.8-94.4) for normal and tumor samples, respectively.

### Somatic mutations in the Discovery Cohort

After filtering, we observed a median mutation burden of 1.31 mutations/Mb (range: 0.033-3.28), comprised of 2,633 SNVs and INDELs across all samples (range: 2-200, median: 77.5, mean=87.8) (**Figure 1**, **Table S1**). These variants were discovered across 2,245 genes with 258 of these genes mutated in more than one sample. Using WGS data from the 26 non-cirrhotic samples, we identified 6 genes that were significantly mutated above background mutation rates according to MuSiC: *ALB, APOB, CTNNB1, TP53, RB1*, and *RPS6KA3* (**Figure 1**, **Table S1**). With regards to all methods of sequencing (WGS, RNAseq, CAP1, and CAP2), the most frequently encountered variant was a SNV in the telomerase reverse transcriptase (*TERT*) promoter (C228T; G1295228A), which was identified in 17/30 samples and resulted in overexpression of *TERT* (**Figure S2, Table S2**). Within the exome, *TP53* was the most recurrently mutated gene and was observed in 8/29 of samples (**Table S1**). Beta catenin 1 (*CTNNB1*) was also significantly mutated within this cohort (6/29), whereby the majority of variants occurred at amino acids S37 and S45, both of which reside in a putative GSK3B phosphorylation site in exon 3 (ENST00000349496) (**Figure S3**) [35]. Frameshift mutations in *APOB* were observed in 4/29 of samples (**Table S1**). Mutation signatures using the COSMIC database for the discovery cohort were investigated. Signatures 5 (unknown etiology), 4 (smoking damage association), 16 (unknown etiology), and 12 (liver damage association) were most prevalent and contributed to the overall cohort signature at 23%, 14%, 8%, and 7%, respectively (**Figure S4**).

**Figure 1:**
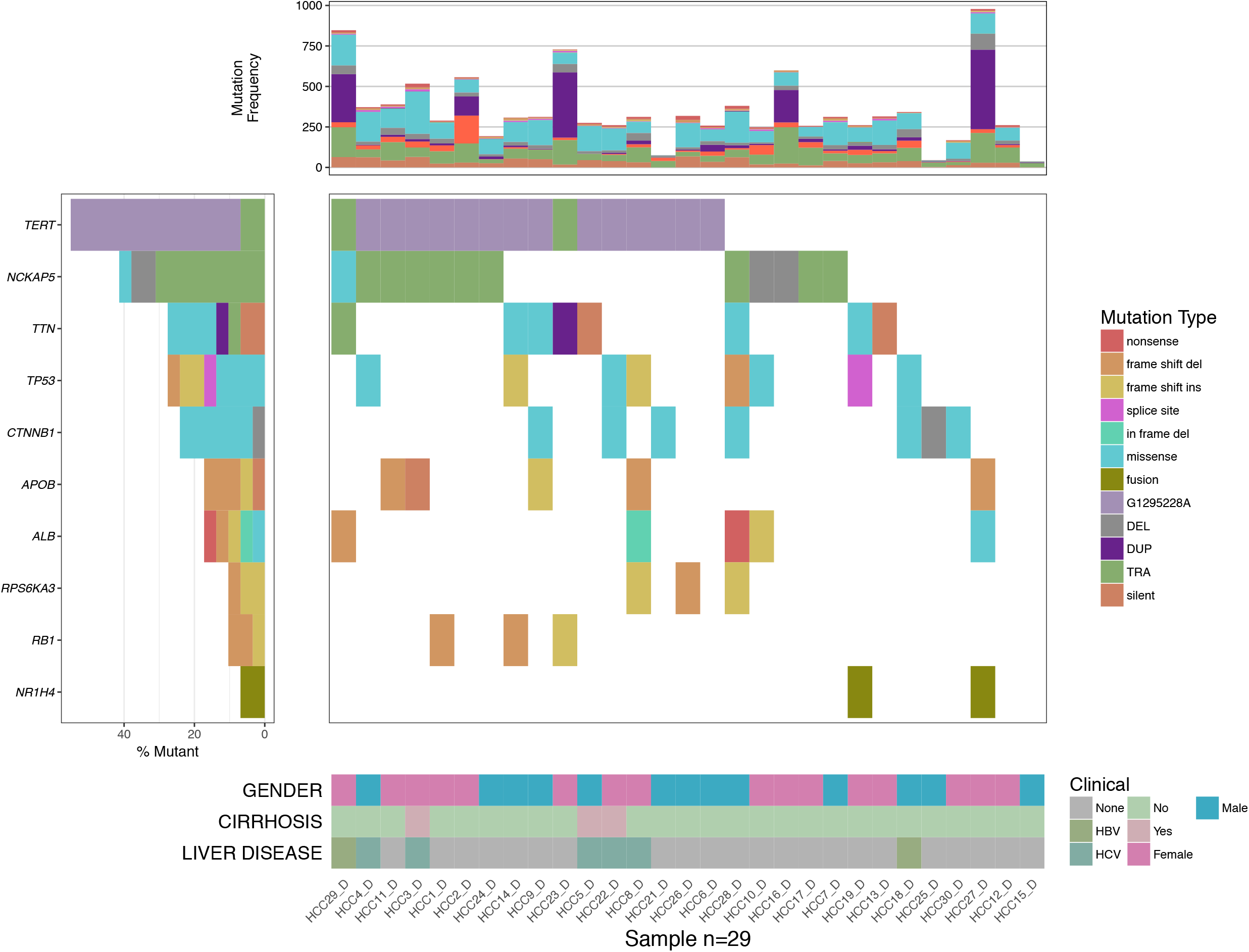
Genomic landscape of the non-cirrhotic discovery cohort exhibits similarity with cirrhotic HCC. GenVisR [80,81] depiction of coding somatic mutations, structural variants, *TERT* promoter mutations (G1295228A), and validated fusions are shown for samples in the discovery cohort which were recurrently (>25%) or significantly mutated. Where there are multiple mutations for the same gene/sample, the most severe mutation is displayed (severity follows the order listed in the legend). The percentage of samples for which a gene is mutated is shown on the left. Mutation Frequency represents the total number of mutations within individual samples. DUP=duplication; DEL=deletion; TRA=translocation.

### Transcriptome Analysis of the Discovery Cohort

Differential gene expression analysis performed on the non-cirrhotic samples revealed that 11% of genes, including *TERT*, were upregulated (4,468/39,392) and 10% of genes, including *CTNNB1* and WISP2, were downregulated (4,114/39,392) compared to adjacent non-tumor liver tissue (q-value < 0.1) (**Table S1**). Comparison of gene log2 fold changes derived from the differential expression analysis revealed the cell cycle pathway as upregulated in the KEGG signaling and metabolism database (q-value ≤ 0.05). Similarly, we observed 16 pathways as down-regulated (q-value ≤ 0.05), most of which are related to metabolic liver processes. Genes such as *ADH5* and *EHHADH* were observed with reduced expression levels and participate in 38% (6/16) of these pathways. Using the Gene Ontology biological process database, we observed 107 pathways as significantly upregulated (q-value ≤ 0.05). The majority of the upregulated pathways were related to cellular division and DNA repair. In addition, 28 pathways were identified as significantly downregulated (q-value ≤ 0.05), many of which were related to liver metabolism (**Table S2**).

### Telomere lengths in the Discovery Cohort

When evaluating the samples within the discovery cohort for telomere length at the DNA level, we observed that the majority of tumor samples exhibited shortened telomeres compared to their paired normal sample (p-value = 0.00011) (**Figure S2**). One exception was seen in sample HCC16_D, which was distinguished by abnormally high expression of *TERT* (FPKM=36) (**Figure S5**).

### Copy Number Variants and Loss of Heterozygosity in the Discovery Cohort

We observed recurrent large scale amplification of the q-arm of chromosome 1 in ≥ 50% of the discovery cohort. Similarly, large scale deletions of the p-arms of chromosomes 8 and 17 were found in ≥ 40% of the cohort (**Figure 2**). In total, analysis with GISTIC and subsequent manual review revealed 75 unique regions across 17 chromosomes as recurrently amplified and 45 unique regions across 17 chromosomes as significantly deleted (q < 0.05) (**Table S1**). No significant associations with tumor differentiation status were made (α=0.05). Each CNV and LOH event was tested for their association with overall survival and recurrence free survival but no significant association could be made following multiple test correction. A total of 33 genes identified as recurrently deleted by GISTIC showed concordant decreased expression in tumor samples (**Table S1**). These include genes previously characterized as relevant to HCC development and progression: *HEYL* [36] (q-value = 0.032), *UQCRH* [37] (q-value = 0.032), and *MUTYH* [38] (q-value = 0.048). A subset of these genes have also been implicated in tumorigenesis, metastasis, and progression of other cancer types and may prove to be relevant for HCC development and progression: *RPL11* [39] (q-value = 0.048), *UBE2D3* [40] (q-value = 0.032), *ARRB1* [41] (q-value = 0.032), *ENG* [42] (q-value = 0.049), and *ABLIM2* [43] (q-value = 0.032).

**Figure 2:**
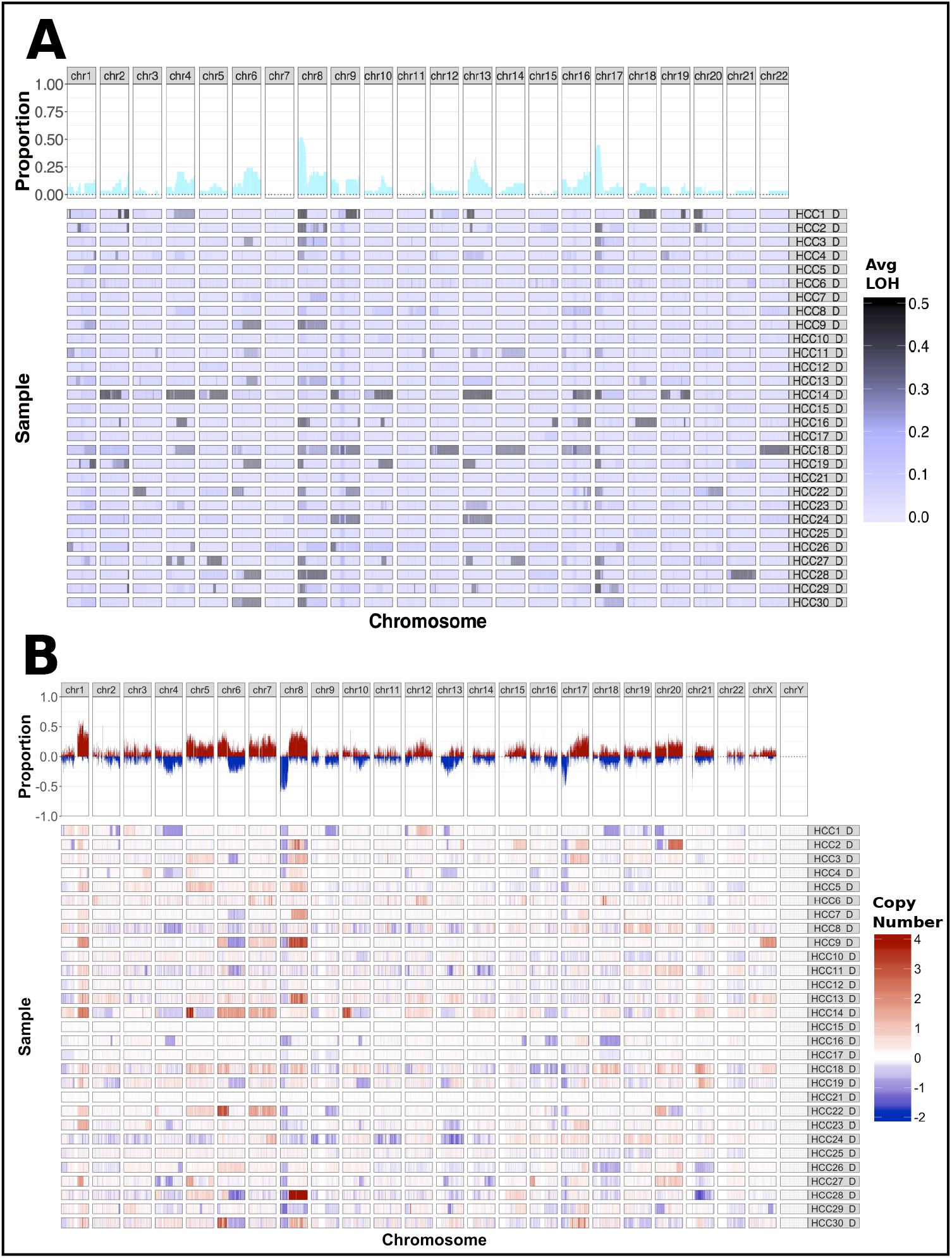
Genome wide CNV and LOH in the discovery cohort. Recurrent regions of LOH (**A**) and CNV amplification/deletion (**B**) are shown for samples within the discovery cohort using the Bioconductor package, GenVisR [80,81]. The proportion of samples with LOH (dark blue), copy number gain (red), and copy number loss (blue) within each chromosomal region are depicted at the top of the panel.

A GISTIC analysis coupled with manual review identified 8 unique regions exhibiting recurrent LOH affecting the chromosomal arms of 6q, 8p, 13q, and 17p (**Figure 2**, **Table S1**). The 8p and 17p chromosomal arms were most susceptible to LOH, each occurring in ≥ 30% of samples. The 8 genomic regions identified as recurrently affected by LOH contain the coding regions for: *TP53, RB1, DLC1, PFN1, ARID1B, LAMA2*, and *CLU* (**Table S1**).

### Structural Variation in the Discovery and Extension Cohorts

We identified 4,745 SV events affecting 3,801 genes across the discovery cohort, of which 737 were deletions, 1,650 were duplications, 450 were inversions, and 1,908 were translocations. Three genes were affected by SVs in ≥ 20% of samples, *NCKAP5* (N=11), *WDPCP* (N=6), and *MACROD2* (N=6) (**Table S1**). Translocations near the *TERT* promoter region occurred in 2/29 samples, both of which were non-cirrhotic (**Figure S5**). Additionally, we detected a recurrent fusion involving *NR1H4* with a diverse set of gene partners (173 fusion predictions) (**Table S1**). NanoString nCounter Elements^tm^ Technology was used to assess 21 of these fusions. NanoString validated three *NR1H4* fusions that were partnered with *EWSR1, GNPTAB*, and *FNIP1* in 2 samples (**Figure 3**). One fusion (*CDK17-NR1H4*) was called by both Integrate and ChimeraScan but was not validated using NanoString. BLAT alignments revealed that these fusions result in sequence frameshifts and thereby likely inactivate the function of *NR1H4*.

**Figure 3:**
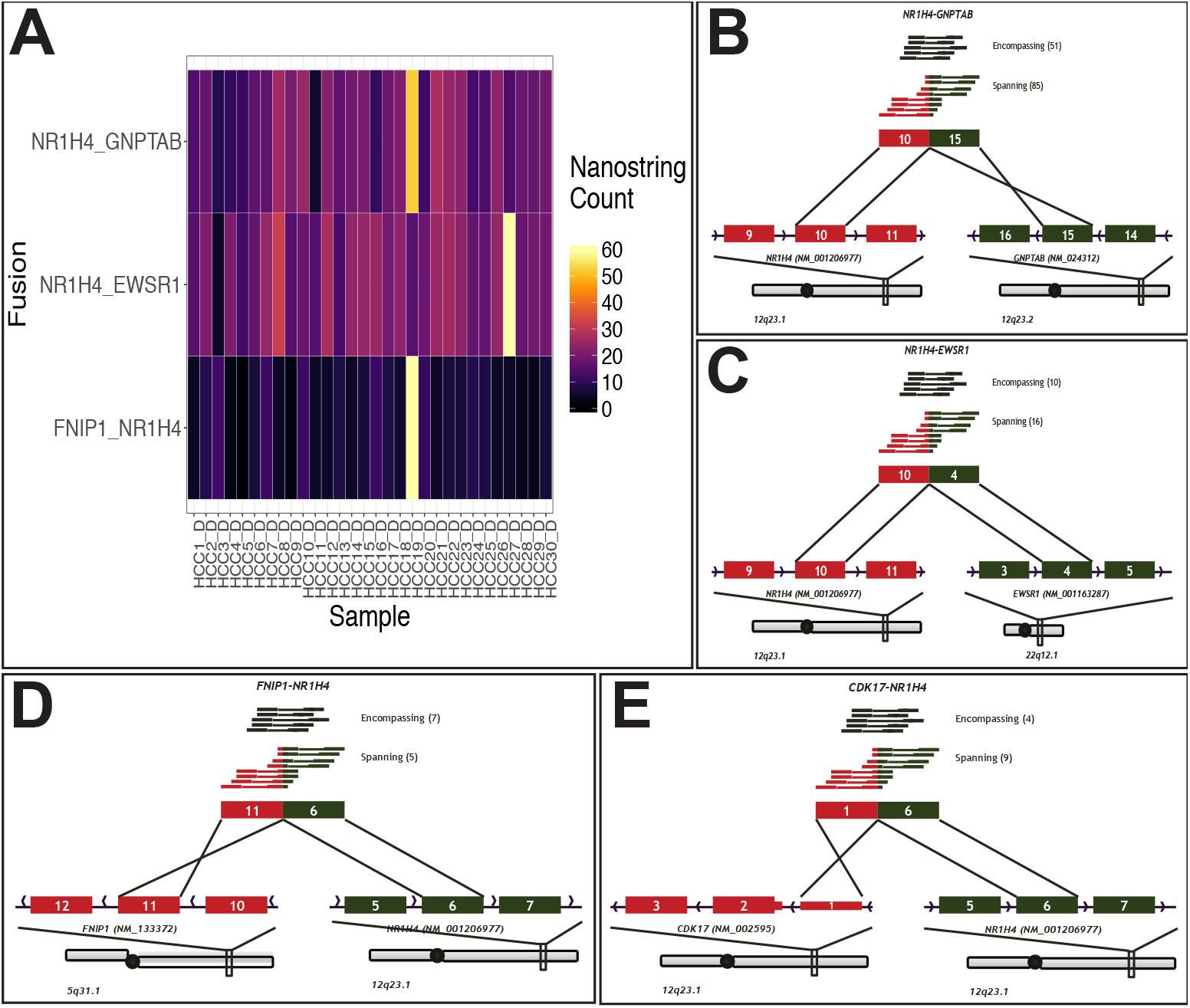
NanoString validation of *NR1H4* fusions observed in the discovery cohort. **A.** NanoString count values for fusions *FNIP1-NR1H4, NR1H4-EWSR1*, and *NR1H4-GNPTAB* across samples in the discovery cohort. Fusion diagrams are shown for: (**B**) *NR1H4-GNPTAB*, (**C**) *NR1H4-EWSR1*, and (**D**) *FNIP1-NR1H4*. **E.**The *CDK17-NR1H4* fusion event was detected by Integrate and ChimeraScan but not validated with NanoString.

### Variant Detection in Extension Cohorts

To further study recurrently mutated genes and discover novel events intrinsic to non-cirrhotic HCC, we employed CAP1 on extension-alpha and extension-beta samples (N=87). In addition to variants identified in the discovery cohort, this extension study elucidated 69 and 1,022 variants in the extension-alpha and extension-beta cohorts, respectively (**Figure S6**). Using the variants identified with CAP1, we classified a total of 17 genes as significantly mutated (q-value ≤ 0.05) (**Table S1**). Of the significantly mutated genes in the discovery cohort, all were confirmed as significantly mutated in the extension cohorts with the exception of *RB1* and *RPS6KA3*. We tested for differences between mutated genes based on cirrhosis status within all cohorts using CAP1, but we were unable to identify any significant differences (q-value ≤ 0.05).

### Germline mutations in Discovery and Extension-Alpha Cohorts

Within the discovery cohort and extension-alpha cohorts, there were 4 genes recurrently mutated (≥4 tumors) in germline DNA after filtering, including: *AL356585.1, MUC19, SVIL*, and *DNAH5*. In addition, when evaluating deleterious calls predicted by four different methods (SIFT [44], Polyphen [45], ClinVar [46], CADD [47]), there were 11 variants that were identified as pathogenic. Of these 11 variants, 5 were in autosomal genes (*LAMA2*, *CYP4V2*, *SLC22A5*, *BMPR2*, *SLC26A4*) and 6 were in mitochondrial genes (*MT-CO1*, *MT-CO1*, *MT-ND3*, *MT-ND3*, *MT-CYB*, *MT-ND1*) (**Table S1**).

### Viral Integration and *TERT* Promoter Mutation in Discovery and Extension Cohorts

Viral detection of HBV and HCV in samples from the discovery and extension cohorts was conducted using CAP2. This analysis validated the clinical diagnosis of HBV in 7 of 9 HBV+ HCC samples. Additionally, a clinically undiagnosed sample in the discovery cohort (HCC18_D) was shown to possess HBV infection both at the RNA and DNA level (**Table S1**). Manual review and BLAT analysis confirmed that HBV integrated at the *TERT* promoter locus in 1 out of the 10 HBV positive samples (**Figure S1**). Similarly HCV, a RNA virus, was detected in 5 samples in the discovery cohort, 3 of which confirmed a clinical HCV diagnosis. Within the extension cohort, 29 samples were clinically diagnosed with HCV. Competitive alignments using kallisto [28] were performed to detect integration of AAV1 and AAV2 in samples from the discovery cohort. AAV1 and AAV2 were both detected in one sample. AAV2 was detected in two other samples (**Table S1**).

Mutations in the *TERT* promoter region were detected in samples in the discovery and extension cohorts with WGS, Sanger sequencing, and the CAP2 panel using Nimblegen and spiked-in IDT probes. Point mutations in this region (C228T or C250T) were observed in 52.4% of the samples. Samples infected with HCV, determined by clinical assay, were found to be significantly enriched for *TERT* promoter mutations (p-value = 0.0051, **Table S1**); however this was no longer significant following a multiple test correction (q>0.05).

### Clinical Associations and Survival Analysis

Among non-cirrhotic HCC samples in the discovery cohort, *TERT* promoter alterations (point mutations, HBV integration, and translocations) were not significantly associated with overall survival or recurrence free survival (**Figure 4**). No significant associations were observed between any other variants (SNV/INDEL, CNV, LOH, and SV) and clinical variables including lymphovascular space invasion, tumor differentiation, and tumor predisposition (e.g. HBV/HCV infection, alcohol abuse, cirrhosis, etc.).

**Figure 4:**
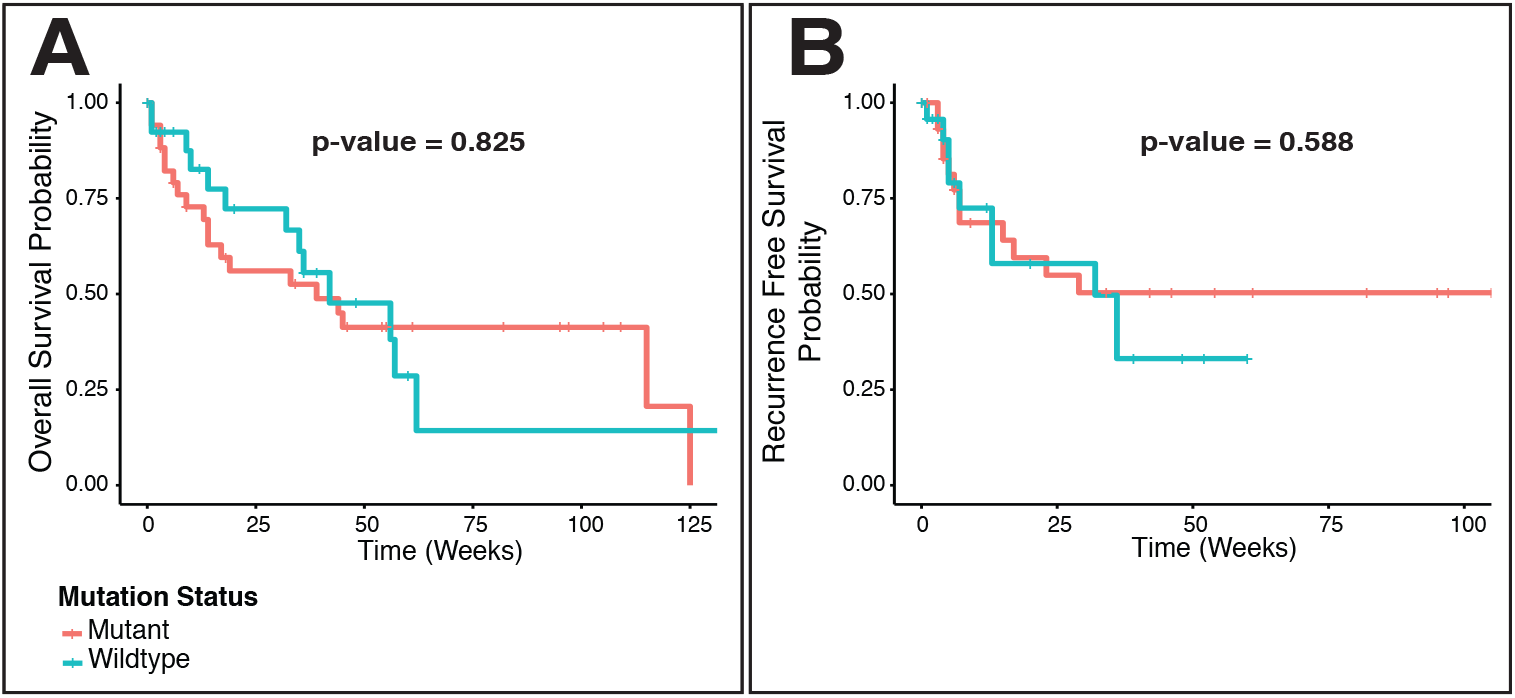
Overall and recurrence free survival analysis for *TERT* promoter mutations. Kaplan Meier curves for *TERT* mutation status (C228T/C250T promoter mutation, HBV promoter integration or *TERT* structural variant) for all non-cirrhotic samples. The probability of overall survival (OS) (**A**) and recurrence free survival (RFS) (**B**) across all non-cirrhotic samples is shown.

## Discussion

### Genomic Landscape of Non-cirrhotic HCC Largely Resembles that of Cirrhotic HCC

The overarching purpose of this study was to test if genomic differences exist between non-cirrhotic HCC and HCC developed in the background of cirrhosis. Within our non-cirrhotic discovery cohort of 26 patients sequenced by WGS, the median mutation burden and recurrent somatic/germline mutations closely resembled those previously reported in cirrhotic HCC [48]. Recurrently mutated genes in the discovery cohort included *TERT* (55%), *TP53* (28%), *CTNNB1* (21%), and *APOB* (13%), all of which have been previously observed in cirrhotic HCC at similar frequencies [9,48]. We did not find any significant difference when comparing recurrently mutated genes between 52 cirrhotic and 63 non-cirrhotic samples for which cirrhotic status was available using CAP1. The LOH events [10,49–52] and CNVs [9–11,52,53] observed within our cohort are similar to those previously observed in cirrhotic HCC using alternative, investigative approaches including SNP array analysis. Within these regions, several genes such as *LAMA2, ATK3, EFF1A1*, and *PFN1* [14,54–56] have been previously investigated in the context of HCC development and progression.

Structural variants identified in non-cirrhotic patients’ samples were also largely similar to SVs reported in cirrhotic HCC. However, of the three genes that were mutated in over 20% of samples in the discovery cohort, only *MACROD2* had been previously described. With its ADP-ribosyl-hydrolase activity, *MACROD2* has a well characterized role in reversing ADP-ribosylation of proteins involved in a variety of cellular processes, including the DNA repair pathway [57]. A study in Japan identified 32/268 HCC samples (cirrhosis status not specified) with SVs involving *MACROD2*, which was associated with a better overall prognosis [8]. Though our data did not support this same clinical association, our findings suggest a similar mutation frequency of *MACROD2* in non-cirrhotic HCC. The observed *TERT* structural variants, which occurred in 2 non-cirrhotic samples, have also been detected in cirrhotic HCC. This study corroborates previous findings of *TERT* activation via SVs, HBV integration, and point mutations in HCC patients [8,9].

Several studies have associated accumulation of β-catenin with cirrhotic HCC tumorigenesis [58]. However, in cases of HCC with reduced inflammation and fibrosis, researchers have observed an absence of β-catenin accumulation [59]. It is therefore intriguing that despite observing activating mutations within exon 3 of *CTNNB1*, we observe *CTNNB1* downregulation in non-cirrhotic samples of the discovery cohort. Accumulation of β-catenin is possible in the context of *CTNNB1* down-regulation since exon 3 mutations serve to prevent the degradation of the β-catenin protein [58]. The majority of other studies observe accumulation of β-catenin in the context of a primarily asian cohort whereas our cohort represents a western population. Future investigations are required to understand the role of *CTNNB1* expression and β-catenin accumulation in non-cirrhotic HCC.

### Biological Pathways in Liver are Dysregulated

Investigations into significantly enriched pathways in non-cirrhotic samples from the discovery cohort revealed broad downregulation of pathways related to liver metabolism (GO [60,61], KEGG [62–64]) and upregulation of pathways involved in cellular division and replication (GO) (**Table S2**). This is expected given that tumor cells are dividing more frequently and losing normal liver function. We observed downregulation of pathways linked to cytochrome P450 (CYP450) mediated xenobiotic metabolism. Previous investigations of CYP activity and expression in cirrhotic and HBV infected HCC demonstrate that CYP activity is dysregulated in HCC tumor cells [65]. Given the potential for CYPs to facilitate individualized treatment options for HCC patients, it is possible treatment strategies for non-cirrhotic HCC may also involve CYPs.

### Survival Analysis Does Not Identify Prognostic Potential for Observed Mutations

A survival analysis was performed on non-cirrhotic samples of the discovery cohort to identify large scale mutational events (SVs, CNVs, and LOH) that may serve as prognostic biomarkers; however, no significant association could be made. The survival analysis was also extended to SNVs and INDELs. *TERT* promoter alterations have been associated with poor prognosis in cancers such as glioblastomas [15,66] and melanomas [16]. In terms of HCC however, previous reports that investigated *TERT* promoter status with respect to survival yield conflicting results [23,67]. We found no association between *TERT* promoter mutations and prognosis. Taken as a whole, it appears activating *TERT* mutations serve as a common mechanism for tumorigenesis but additional investigations are required to definitively determine whether or not these mutations can serve a prognostic role in patients afflicted with HCC.

### Low-frequency, Novel Mutations in *APOB* and *NR1H4* and Novel Structural Variants Identified

In this study, we identified predicted loss of function/damaging somatic and germline *APOB* mutations in 6/26 non-cirrhotic samples in the discovery cohort (**Table S1**). Among the somatic *APOB* mutations, all were observed at a relatively high VAF (>0.25), suggesting these mutations were present early in tumorigenesis and may be present in pre-malignant sites within these tissues. It has been shown that abnormal glycosylation resulting from the formation of bisecting-GlcNAc disrupts APOB function and leads to a fatty liver disease phenotype [68]. We observed *APOB* mutations outside of a fatty liver disease phenotype suggesting a more prominent role in tumorigenesis.

We also identified a novel recurrent somatic, loss of function, gene fusion event involving *NR1H4* in 2 non-cirrhotic samples within the discovery cohort (**Table S1**). Low sample quality (particularly FFPE samples) and use of RNA-based validation methods may have prevented sensitive detection of these fusion events resulting in underestimation of fusion frequency in additional cases. Biologically, *NR1H4* has a well-defined role to prevent the accumulation of bile acid (BA) within the liver, which could otherwise lead to HCC development. *In vivo* studies have demonstrated that *NR1H4* loss predisposes mice to spontaneous hepatocarcinogenesis [69,70] and obstructs hepatocyte regeneration following partial hepatectomy [71]. *NR1H4* fusions may therefore represent a novel mechanism of HCC development.

Other novel SV events that were observed in the discovery cohort involved *NCKAP5* and *WDPCP*, both of which play a role in cilia function. *NCKAP5L*, which is a paralog to NCKAP5, functions to stabilize and strengthen microtubule structure [72]. Additionally, reduced expression of *WDPCP* has been shown to inhibit proper ciliogenesis in the presence of proinflammatory cytokines [73]. Given the previous association of ciliopathies with cancer, these recurrent structural variants in non-cirrhotic HCC might play a role in tumorigenesis and metastasis [74].

### Genotoxic and virologic etiologies partially explain Non-Cirrhotic HCC

Recent work by *Zucman-Rossi et al*. has outlined etiologies for HCC development [75]. Within our discovery cohort, 23/29 samples exhibited a mutational signature of T->C mutations at an ApTpN context (weight > 0.1) (**Table 2, Figure S4**). These signatures (signature 5,16) have been associated with genotoxic injury and were previously observed in HCC patients with high alcohol and tobacco consumption. We do not have data on tobacco consumption; however, most of these cases did not have reported alcohol consumption (17/23), which suggests that another unknown factor may be contributing to this observation. Among these 23 samples, a non-cirrhotic case exhibited a signature consistent with aflatoxin exposure (signature 24) and harbored an R249S mutation in *TP53*. Interestingly this case was clinically diagnosed with HBV, which has been suggested to have a synergistic effect with aflatoxin exposure, facilitating HCC development [75,76]. In addition, we observed a single HCV positive case which also possessed signatures 5 and 16. It is curious that while HCV is a predisposing factor for liver cirrhosis, and this patient exhibited cirrhosis, etiologies associated with signatures 5 and 16 are not typically associated with cirrhosis [75]. With the exception of a final HBV case, we could not identify an etiology for the remaining 5 samples. The dysregulation of liver metabolism identified in the pathway analysis may represent a cause or symptom of the etiologies leading to tumorigenesis. This could apply to the unknown etiologies in the remaining samples, suggesting liver dysfunction is necessary for disease progression. Emerging evidence suggests that HCC can develop in the context of nonalcoholic fatty liver disease (NAFLD) or NASH, even in the absence of advanced fibrosis or cirrhosis [77–79]. Our understanding and recognition of the clinical features associated with NASH and NAFLD has evolved. Therefore, it is likely that patients were affected by these conditions, but not clinically diagnosed at the time, within our discovery cohort. Further research is needed to elucidate the relationship between etiologies of HCC and their association with liver metabolism.

## Conclusion

It has been observed that the underlying etiologies contributing to tumorigenesis of non-cirrhotic and cirrhotic HCC are unique [75]. Despite distinct evolutions of these tumor subtypes, our findings describe a convergence of both subtypes onto a similar genomic landscape during disease progression. This genomic similarity suggests in vitro and in vivo models for investigating HCC biology may be relevant to both HCC subtypes for advanced disease. Clinically, genomic-based diagnostic, prognostic, and treatment strategies that were previously established in patients with cirrhotic disease may also be extended to patients with progressed non-cirrhotic HCC.

## Supporting information

List of tables and figures

---

**Figure S1:**
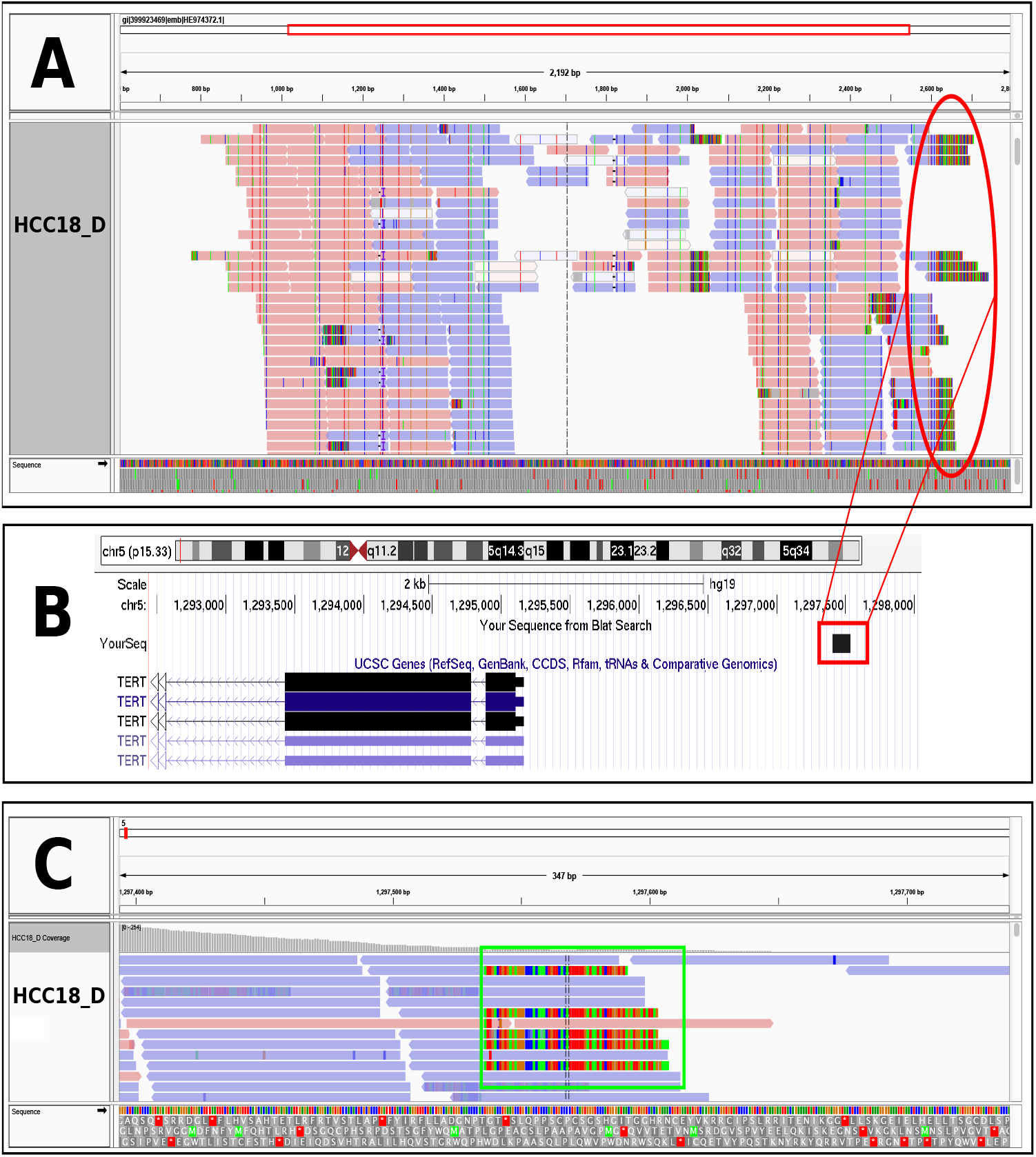
Evidence for a case of undiagnosed HBV integration at the *TERT* promoter. **A.** WGS reads originating from a competitive alignment between GRCh37 and HBV are shown for HCC18_D using the Integrative Genomics Viewer (IGV). **B.** The soft clipped sequence from panel A (red box) was inputted into BLAT to show that it aligns upstream of the *TERT* transcription start site (TSS). **C.** Reads which are soft clipped in panel A are shown aligning to the intronic region of human chromosome 5 in the HCC18_D tumor sample upstream of the *TERT* TSS.

**Figure S2:**
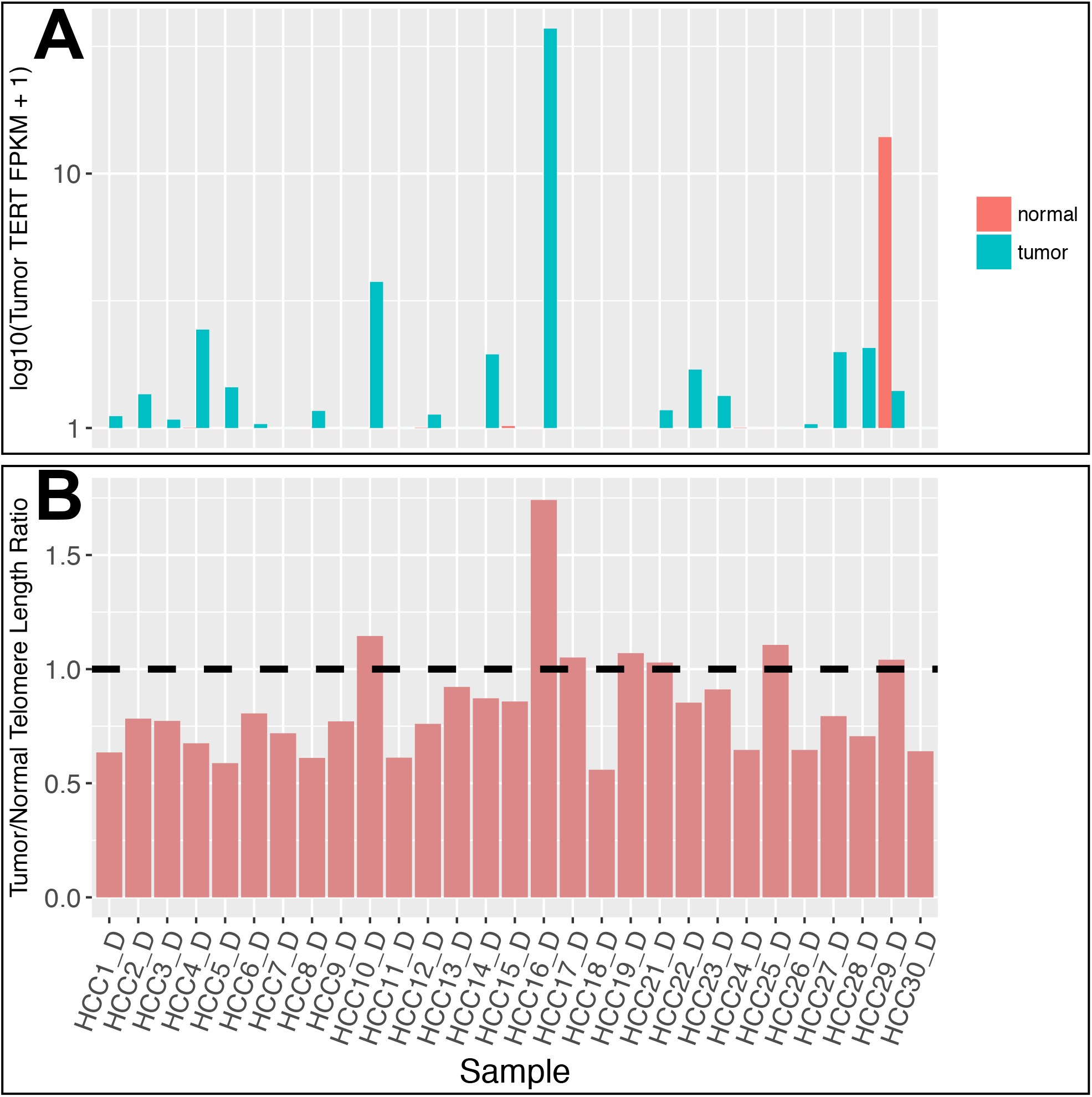
HCC tumors exhibit shortened telomeres. **A.** *TERT* expression (log10(FPKM + 1)) within each matched tumor and normal sample from the discovery cohort, excluding HCC20 (top panel). **B.** Tumor:normal read ratios of telomeric regions are plotted as vertical bars in the bottom panel. The black horizontal line represents a read ratio of 1.

**Figure S3:**
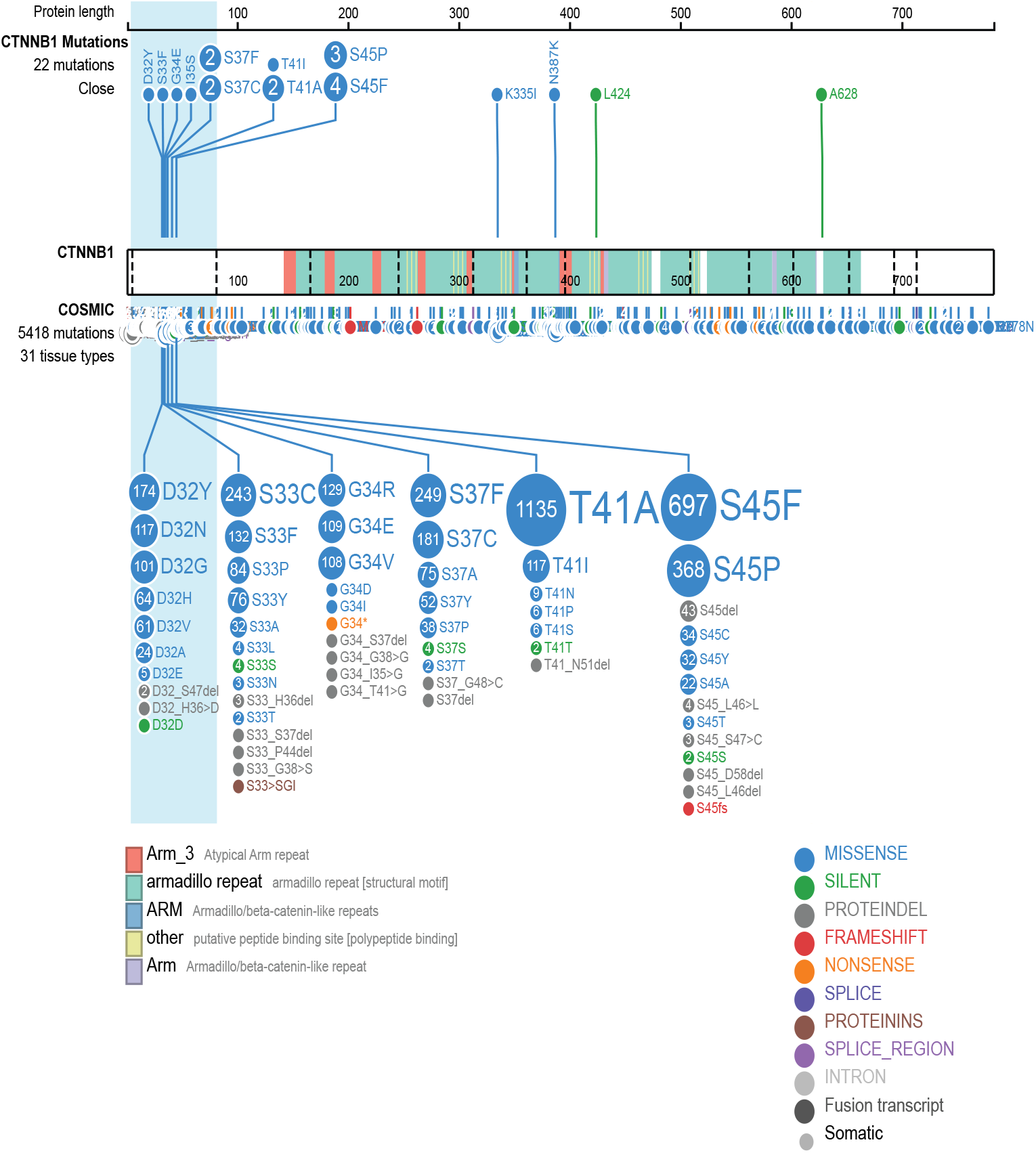
GSK3B phosphorylation site mutational hotspot in *CTNNB1* is observed in non-cirrhotic HCC. ProteinPaint [78] diagram illustrating amino acid (AA) changes within the coding sequence of *CTNNB1*. Legends describing the functional domains and variant types of the AA changes along *CTNNB1* from our dataset and that of COSMIC’s database are indicated at the bottom of the figure. The light-blue highlighted region indicates exon 3 of the *CTNNB1* gene.

**Figure S4:**
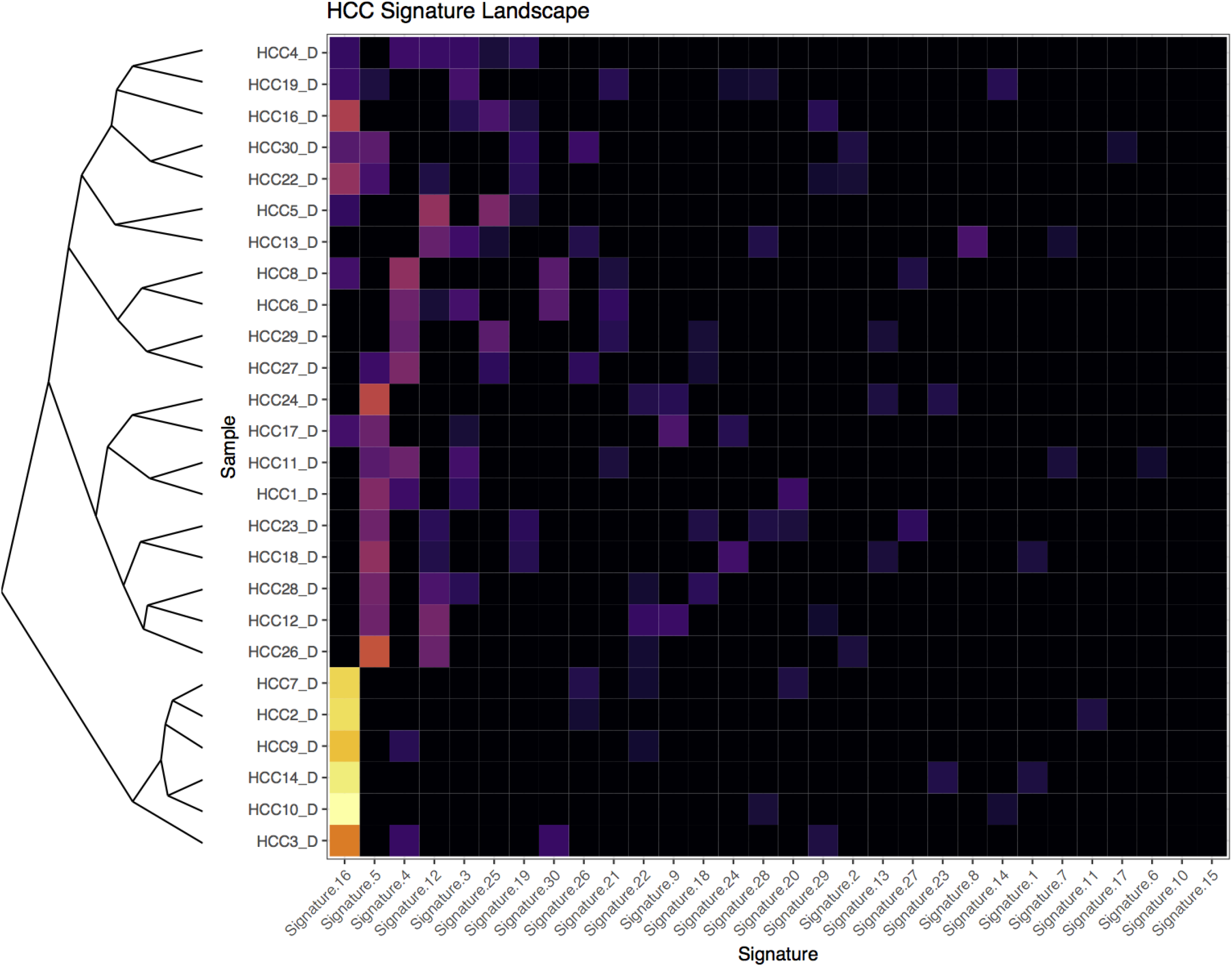
Discovery cohort samples are primarily composed of 4 mutational signatures. The weights from 0 to 1 of relative COSMIC mutational signature distribution are shown for each sample in the discovery cohort. Samples are hierarchically clustered using Euclidean distances based on assigned weights, signatures are ordered from left to right by decreasing weight.

**Figure S5:**
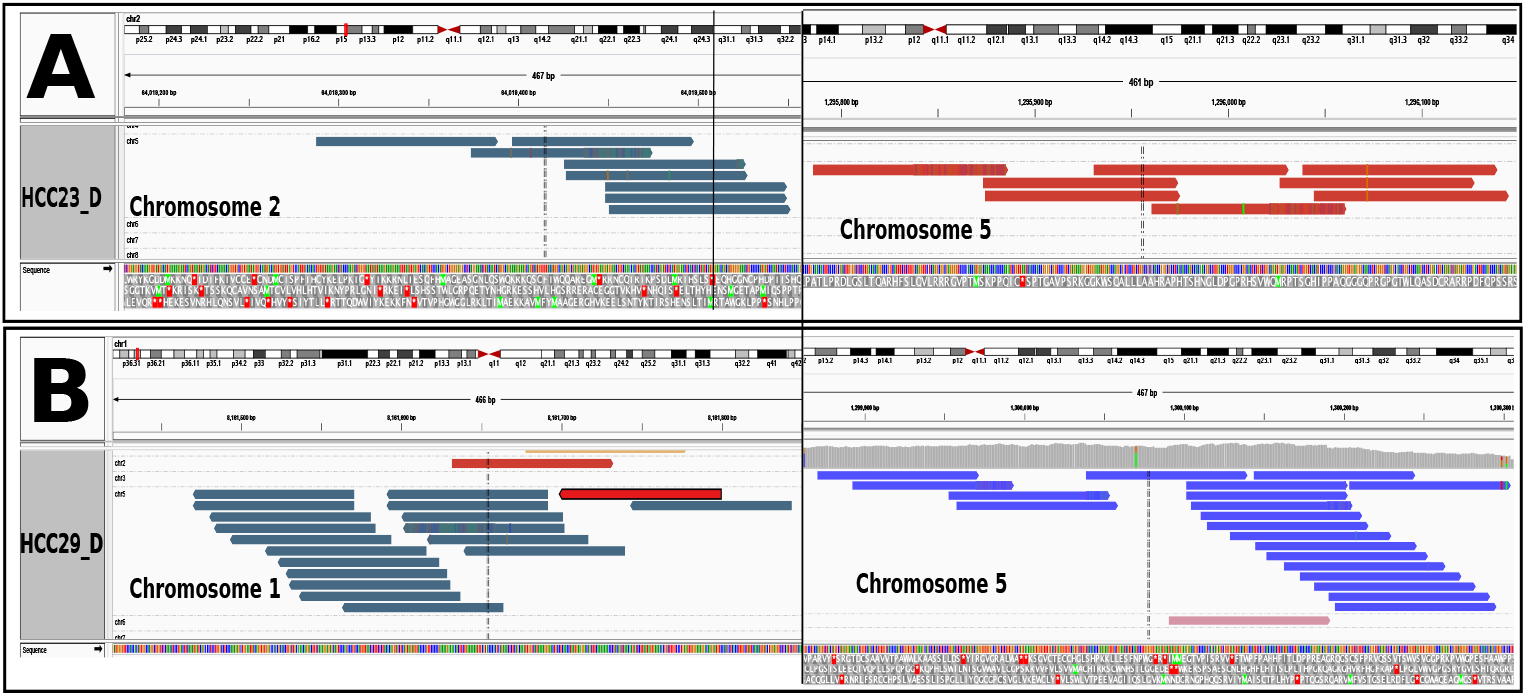
Evidence for translocations involving *TERT*. Reads depicting translocation events involving *TERT* in tumor samples for (**A**) HCC23_D and (**B**) HCC29_D are shown. In both figures, the left panels contain dark blue reads that align to regions of chromosome 2 (HCC23_D) and chromosome 1 (HCC29_D). Their paired read aligns to a region ~2-3kbp upstream of the *TERT* promoter on chromosome 5, shown on the right.

**Figure S6:**
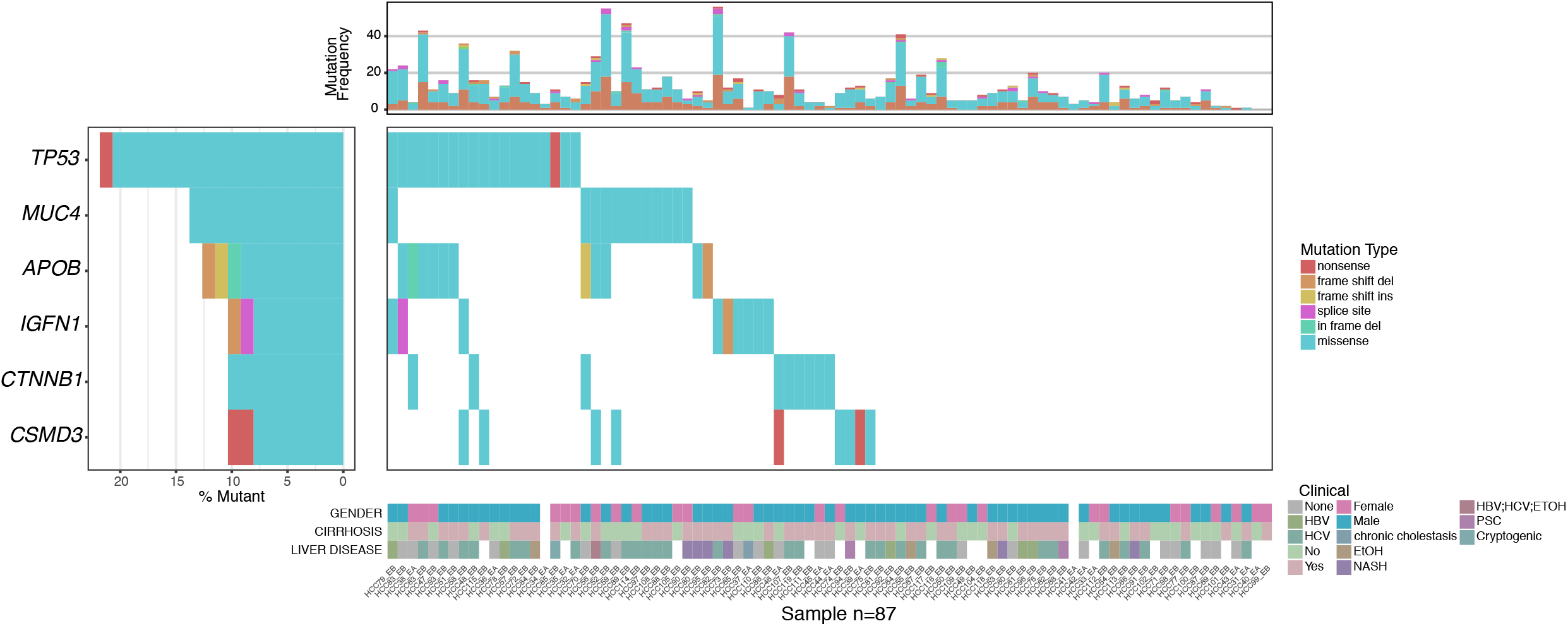
Genomic landscape of the extension cohort. Manually reviewed coding, somatic mutations from the extension alpha and beta cohorts (N=87) derived from CAP1. In situations where there are multiple mutations for the same gene/sample, the most deleterious mutation is shown following the order of the mutation type legend. Clinical variables for each sample are included at the bottom. Tumor mutational frequency for each sample is plotted at the top.

