## Supplementary material for "Genomic and transcriptomic somatic alterations of hepatocellular carcinoma in non-cirrhotic livers": List of tables and figures

### Main Tables

**Table 1 - Cohort Summary**
**Table 2 - Proposed etiologies**

### Main Figures

**Figure 1 - Genomic landscape of the non-cirrhotic discovery cohort exhibits similarity with cirrhotic HCC**
**Figure 2 - Genome wide CNV and LOH in the discovery cohort**
**Figure 3 - NanoString validation of *NR1H4* fusions observed in the discovery cohort**
**Figure 4 - Overall and recurrence free survival analysis on *TERT* promoter mutations**

### Supplemental Tables

**Table S1: Sample details for discovery and both extension sets**
**Table S1: Clinical information for samples in discovery and both extension cohorts**
**Table S1: Somatic variants found in discovery and both extension cohorts**
**Table S1: SMG list for all samples in discovery and both extension cohorts**
**Table S1: Germline variants observed in discovery and extension-alpha cohort**
**Table S1: Fusions called by ChimeraScan and Integrate for discovery samples**
**Table S1: Genes affected by Manta-reported structural variation in discovery cohort**
**Table S1: Downregulated genes in all samples from discovery cohort**
**Table S1: GISTIC reported recurrent regions of CNV and LOH in the discovery cohort**
**Table S1: Genes with copy number loss and concordant decrease in expression**
**Table S1: HBV/HCV/AAV1/AAV2 status for samples in discovery cohort following sequencing at the DNA and RNA level**
**Table S1: TERT mutations observed in the discovery and extension cohorts**
**Table S2: GAGE Pathway Analysis**
**Table S3: Capture Panel I Design**

### Supplemental Figures

**Figure S1: Evidence for a case of undiagnosed HBV integration at the *TERT* promoter**
**Figure S2: HCC tumors exhibit shortened telomeres**
**Figure S3: GSK3B phosphorylation site mutational hotspot in *CTNNB1* is observed in non-cirrhotic HCC**
**Figure S4: Discovery cohort samples are primarily composed of 4 mutational signatures**
**Figure S5: Evidence for translocations involving *TERT***
**Figure S6: Genomic landscape of the extension cohort**
