## Supplemental Methods for "Genomic and transcriptomic somatic alterations of hepatocellular carcinoma in non-cirrhotic livers"

### Supplementary Methods

#### Sample Acquisition

The discovery cohort consisted of surgical resections for 30 primary tumors, whereby matched adjacent non-tumor liver tissue comprised the normal controls. These samples did not develop in the context of hepatocellular adenoma (HCA) and were banked between 2000 and 2011 from patients diagnosed with HCC at the Washington University School of Medicine. The extension cohort was comprised of an additional 87 primary tumors, wherein 16 HCC tumors (with matched non-tumor liver) were used for extension-alpha and 71 HCC tumors (no matched normals) were used for extension-beta (**Table 1**). All tissue samples for the discovery and extension-alpha cohorts were flash-frozen prior to banking, extension-beta tissue was derived from formalin fixed paraffin embedded (FFPE) blocks. All patient samples were acquired after informed consent to an approved study by the Washington University School of Medicine Institutional Review Board (IRB 201106388) (**Table 1**). Across the three cohorts, 73 were male and 44 were female. Additionally, 15 were African American, 98 were Caucasian, and 2 were Asian.

#### Sample Preparation and Sequencing

DNA and RNA were extracted from the discovery cohort samples using QIAamp DNA Mini kit and Qiagen RNeasy mini kit respectively. Following these isolations, we evaluated the quality and quantity of DNA and RNA prior to Illumina library construction. Genomic DNA was sheared by Covaris, size-selected, and Kapa HYPER kits were used to construct libraries for whole genome sequencing by Illumina. We aimed for 60X coverage for tumor samples and 30X coverage for normal samples from the HiSeq 2000 (paired-end 100 bp reads). RNAseq libraries were constructed using the Ovation RNA-seq System V2 (NuGen Inc) kit and the resulting barcoded libraries were pooled prior to Illumina sequencing aimed at 50M reads per library. DNA was extracted using QIAamp DNA Mini kit for extension-alpha. A hybrid capture panel (CAP1) was designed based on preliminary data from WGS of the discovery cohort. The CAP1 targets encompass 12,499,707 base pairs and 68,396 distinct regions (**Table S3**). We used this reagent to capture fragments from the discovery WGS libraries and perform Illumina sequencing to validate previously identified variants. CAP1 was also used to identify variants in the extension-alpha cohort. DNA was extracted from the extension-beta cohort using the QIAamp DNA FFPE Tissue kit and variants were identified using the CAP1 sequencing strategy. A second hybrid capture panel (CAP2) was designed using Nimblegen and spiked-in IDT probes that hybridized to the *TERT* promoter locus (chr5:1,295,163-1,295,663) and the HBV consensus genome. This panel was used to evaluate all 117 tumors. The HBV consensus genome was constructed from 10 HBV genotypes. Sanger sequencing was also performed on the discovery and extension-alpha cohorts targeting the *TERT* promoter locus. cDNA capture sequencing was performed on pooled samples from extension-alpha and extension-beta. All sequencing data were deposited in the Database of Genotypes and Phenotypes (dbGAP; accession phs001106).

#### Sequence Alignment

All data were analyzed using the Genome Modeling System (GMS) [1]. For WGS and CAP1, reads were aligned to GRCh37 using BWA (0.5.9, -t 4 -q 5) [2]. For CAP2 data, reads were competitively aligned against the human reference genome (GRCh37) along with ten HBV genotypes (A1, A2, B, C, D3, D4, E, F2, G, and H) for which complete genomes were available (Accession ids: HE974362.1, HE974364.1, AB602818.1, AB644280.1, HE974377.1, HE974372.1, HE974380.1, HE974366.1, AP007264.1, AB516393.1). RNAseq data were aligned with bowtie/tophat (v2.0.4: --bowtie-version=2.0.0-beta, -p 4) and expression was evaluated with cufflinks (2.0.2: -p 4) [3,4]. All raw RNAseq reads from the discovery cohort were also aligned against the HBV genomes for evidence of HBV expression at the RNA level. The most predominant HBV strain was determined by using relative coverage for competitive alignments. To identify the precise location of the HBV integration site, we realigned the HBV CAP2 reads to GRCh37 and to the predominant HBV strain's genome to identify discordant read pairs. A similar procedure was performed for HCV whereby both WGS and RNAseq reads were aligned against six HCV genotypes (1-6) (NC_004102.1, NC_009823.1, NC_009824.1, NC_009825.1, NC_009826.1, NC_009827.1). The predominant HCV strain was determined using the total read support. Given that HCV has an RNA genome, no alignments or integration sites were expected from the WGS data.

**Germline Variant Calling**

Single nucleotide variant (SNV) and insertion/deletion (INDEL) events for the discovery and extension-alpha cohorts were obtained using the somatic variation pipeline within the GMS. Briefly, variant calls originating from normal alignments were obtained with samtools (0.1.16) and read counts acquired using bam-readcount (0.7, -b 20 -q 20). Variants were filtered to have a variant allele fraction ≥ 0.30 and coverage ≥ 20X, with remaining variants annotated using the Ensembl variant effect predictor (VEP) in combination with the CADD plugin (v88, --per_gene, --everything --plugin CADD) [5,6]. Further filtering removed common SNPs based on ExAC (v0.2) [7], 1000 Genomes [8], or the Exome Sequencing Project [9] (allele frequency ≥ 0.001). Variants with mild consequences in severity, based on the VEP guidelines ("intron_variant" or below), were also removed. Remaining variants were selected if at least two of the following four conditions were met: ClinVar status was "pathogenic" or "likely pathogenic", SIFT prediction was "deleterious" or "deleterious low confidence", PolyPhen prediction was "damaging" or "damaging low probability", or scaled CADD score was ≥ 15. Further, we recognized that *de novo* germline variants are relatively unlikely to show recurrence across 30 unrelated samples. Therefore, to eliminate alignment and sequencing artifacts, we removed any variants which were identified in more than two independent samples.

**Somatic Variant Calling and Analysis**

SNV/INDEL events for all samples were obtained using the somatic variation pipelines within the GMS. Briefly, for the 30 WGS samples an INDEL detection strategy using the union of calls from GATK-somatic-indel (v1.0.5536), Pindel (v0.5: -variant-freq-cutoff=0.2), VarScan2 (v2.2.6), and Strelka (v0.4.6.2: -isSkipDepthFilters=0) was employed with default parameters unless otherwise specified [10–13]. Similarly, a SNV detection strategy using the union of calls from samtools (v0.1.16) intersected with Somatic Sniper (v1.0.2; -F vcf -q 1 -Q 15), VarScan2 (v2.2.6), and Strelka (v0.4.6.2: isSkipDepthFilters=0) was used [12–14]. An identical strategy was employed for the extension-alpha capture (CAP1) with the following exception: Strelka was run with and without the isSkipDepthFilters parameter and the union was taken. For the 71 samples in extension-beta, the somatic variation pipeline was run with an INDEL detection strategy, which used calls from VarScan2 (v2.2.6: --min-coverage 3 --min-var-freq 0.8 --p-value 0.10 --strand-filter 1 --map-quality 10). The SNV detection strategy used the union of calls from samtools (v0.1.16), and VarScan (v2.2.6: --min-coverage 3 --min-var-freq 0.08, --p-value 0.10 --strand-filter 1 --map-quality 10) [12]. Variant calls for all samples were restricted to tier 1 (primarily coding) mutations as described in Mardis et al. 2009 [15]. In addition, we removed variants within a gl-contig, variants within a mitochondrial gene, and variants within ExAC (v0.2: allele frequency ≤ 0.001) [7]. The remaining variants were then quantified using bam-readcount (v0.7; -b 20, -q 20). We removed artifacts or probable germline variants by eliminating SNVs that had greater than three reads in two or more normal samples and any INDELs that had greater than two reads in two or more normal samples. The remaining variants underwent additional filtering using a log-likelihood ratio test as previously described [16]. If a normal sample was not available for the log-likelihood ratio test, the comparator coverage was derived from the associated tumor sample and the comparator variant read count was set to zero. A somatic filter was applied in cases with paired normal samples, which removed variants if any of the following was true for SNVs: normal VAF was ≥ 5% and normal coverage ≥ 20x; tumor VAF ≤ 5% and tumor coverage ≥ 20x; or tumor variant reads ≤ 3. INDELS were filtered in an identical manner with the following exception: if tumor variant reads were ≤ 2, they were removed. Finally, variants within non-protein-coding transcripts were removed and all subsequent variants underwent manual review to ensure quality. Variants were annotated using the GMS based on Ensembl version 74_37 [17].

A Fisher's exact test was used to determine enrichment of mutation rates between cirrhotic and noncirrhotic samples in the extension cohorts for all genes with ≥ 5 variants. SNVs passing manual review in the discovery cohort were analyzed for signature motifs using the bioconductor package deconstructSigs (v1.8.0) for all samples with > 50 variants [18]. Signatures were determined using the COSMIC database via the 'genome2exome' method of normalization within the package. Samples were clustered on signature weights using euclidean distances with a complete linkage method. Significantly mutated genes were detected using MuSiC (v0.4) with an alpha value ≤ 0.05 (**Table S1**) [19]. Multiple testing correction was performed using the false discovery rate (FDR) procedure based on a convolution test.

**Telomere Length Determination**
WGS data from the discovery cohort was used to count the number of telomeric reads in matched tumor-normal samples using the GMS [20]. The tumor:normal read ratio for each sample was determined and visualized with ggplot2. A Wilcoxon-Mann-Whitney test was employed to calculate the significance of differences between telomere length across tumor and normal samples.

**Structural Variant Calling**
Somatic structural variants (SV) for the discovery cohort were called using Manta (v0.29.6) [21]. Manta-reported variants were filtered based on the size of the SV event (<500,000 bp) and subsequently manually reviewed using the integrated genome viewer (IGV) [22]. Genes occurring within a 10kb flank of each SV breakpoint were annotated using the bioconductor package biomaRt (v2.28.0) and ensembl (GRCh37.p13) [17,23].

**NanoString nCounter Elements™ Tagsets: *NR1H4* Fusion Validation**
A NanoString nCounter Elements™ Tagsets assay was performed to validate NR1H4 *in-silica* fusion predictions. Briefly, *NR1H4* fusions called by chimerascan (v0.4.6) [24] and integrate (v0.1d) [25] were obtained from the RNAseq data of the discovery cohort. Fusion calls corresponding to the extension cohorts were obtained from a cDNA capture designed to target *NR1H4* and applied to pooled samples for which RNA was available. A filter was applied requiring *in-silica* fusion predictions to have ≥10 reads of total support (spanning + encompassing) and ≥1 read of spanning support. Sequences corresponding to predicted transcripts for these fusion breakpoints were provided to NanoString for probe design. Fusions were omitted from the final validation design if they lacked a suitable probe. The NanoString assay was then applied to all samples for which RNA was available according to manufacturer recommendations. Assay cartridges were loaded with 100-200ng of RNA.

**Copy Number and LOH Analysis**
Somatic copy number variants (CNV) for the discovery cohort were identified using the GMS as previously described [1]. Copy number calls were segmented into regions using the circular binary segmentation algorithm available in the bioconductor package DNAcopy (v1.46.0). Recurrent regions harboring CNVs were identified with GISTIC (v2.0.22; conf = 0.90, savegene = 1, genegistic = 1, broad = 1, maxseg = 15000, qvt = 0.15) [26]. Loss of heterozygosity (LOH)

was determined by comparing tumor/normal VAF differences from heterozygous calls in the normal samples (normal VAF > 0.4 and < 0.6) originating from VarScan2 (2.2.6) [12]. DNAcopy (v1.46.0) circular binary segmentation was used to generate segments of LOH and GISTIC (v2.0.22) [26] was used to analyze recurrence. However, because GISTIC does not directly conduct a recurrence analysis for LOH calls, LOH events were treated as copy number deletion events. To this end, the absolute difference between the tumor VAF and 0.5 was calculated at each heterozygous variant position. An absolute mean deviation (AMD) value was subsequently obtained for each segment. In accordance to the GISTIC input requirements for copy number recurrence analysis, each AMD value was assigned a discrete value along a continuum between 0.0 to -1.0 such that a value of -0.1 was minimal evidence for a real LOH event. All GISTIC-reported chromosomal regions of CNV and LOH were manually reviewed with the IGV. This analysis was restricted to the autosomes for all samples.

Genes within GISTIC-predicted amplified or deleted regions were intersected with a fragments per kilobase per million (FPKM) differential expression matrix created via cufflinks (v2.0.2; --num-threads 4 --max-bundle-length 10000000) [3,26]. A Wilcoxon-Mann-Whitney test was performed to compare the average FPKM difference for each gene across identified mutant and wildtype tumor samples. A subsequent multiple test correction in R was performed using the FDR method. This identified significant differentially expressed genes (q-value < 0.05) that showed a concordant change in expression relative to GISTIC-reported loss or gain.

**Clinical and Survival Analysis**

Survival analysis was performed to identify genomic alterations significantly associated with overall survival and recurrence free survival. The R packages “survival” and “ggplot2” were used to generate Kaplan-Meier curves illustrating these survival statistics [27,28]. This analysis was performed for genes and genomic regions mutated in at least 4 samples from the WGS data within the noncirrhotic discovery sample cohort (CNV, LOH, and SV). SNV/INDELs were also tested across all noncirrhotic samples from all cohorts. All generated p-values were corrected for multiple comparisons using the FDR methodology (q-values < 0.05). Samples without relevant clinical data were excluded from the analysis. The same procedure was done to test for clinical associations with variables: lymphovascular space invasion (LVSI), tumor differentiation status, and liver disease.

**Differential Expression and Pathway Analysis**

Differential gene expression was evaluated with the bioconductor package DEseq2 (v1.14.1) for the noncirrhotic samples within the discovery cohort [29]. Read counts were obtained for tumor and adjacent matched normal tissues using HTseq-count [30] (v0.5.4p1; --mode intersection-strict, -minqqual 1 --blacklist-alignments-flags 0x0104 --result-version 1), whereby genes lacking reads in all samples were omitted from the analysis. Differential expression analysis, based on a negative binomial distribution, was performed for paired tumors/normals using samples as a blocking factor. A Wald test was used to assess significance and the Benjamini & Hochberg method was used for multiple testing correction. Pathway analysis from resulting log2 expression values was performed via the bioconductor package gage (v2.24.0) with default parameters for both KEGG and Gene Ontology databases [31–36]].

### BIBLIOGRAPHY AND REFERENCES CITED

[19] Dees ND, Zhang Q, Kandoth C, Wendl MC, Schierding W, Koboldt DC, et al. MuSiC:

identifying mutational significance in cancer genomes. Genome Res 2012;22:1589–98.
[20] Parker M, Chen X, Bahrami A, Dalton J, Rusch M, Wu G, et al. Assessing telomeric DNA content in pediatric cancers using whole-genome sequencing data. Genome Biol 2012;13:R113.
[21] Chen X, Schulz-Trieglaff O, Shaw R, Barnes B, Schlesinger F, Källberg M, et al. Manta: rapid detection of structural variants and indels for germline and cancer sequencing applications. Bioinformatics 2016;32:1220–2.
[22] Robinson JT, Thorvaldsdóttir H, Winckler W, Guttman M, Lander ES, Getz G, et al. Integrative genomics viewer. Nat Biotechnol 2011;29:24–6.
[23] Durinck S, Spellman PT, Birney E, Huber W. Mapping identifiers for the integration of genomic datasets with the R/Bioconductor package biomaRt. Nat Protoc 2009;4:1184–91.
[24] Iyer MK, Chinnaiyan AM, Maher CA. ChimeraScan: a tool for identifying chimeric transcription in sequencing data. Bioinformatics 2011;27:2903–4.
[25] Zhang J, White NM, Schmidt HK, Fulton RS, Tomlinson C, Warren WC, et al. INTEGRATE: gene fusion discovery using whole genome and transcriptome data. Genome Res 2016;26:108–18.
[26] Mermel CH, Schumacher SE, Hill B, Meyerson ML, Beroukhim R, Getz G. GISTIC2.0 facilitates sensitive and confident localization of the targets of focal somatic copy-number alteration in human cancers. Genome Biol 2011;12:R41.
[27] Wickham H. Ggplot2. Wiley Interdiscip Rev Comput Stat 2011;3:180–5.
[28] Therneau TM, Grambsch PM. Modeling Survival Data: Extending the Cox Model. Springer Science & Business Media; 2013.
[29] Love MI, Huber W, Anders S. Moderated estimation of fold change and dispersion for RNA-seq data with DESeq2. Genome Biol 2014;15. https://doi.org/10.1186/s13059-014-0550-8.
[30] Anders S, Pyl PT, Huber W. HTSeq--a Python framework to work with high-throughput sequencing data. Bioinformatics 2015;31:166–9.
[31] Luo W, Friedman MS, Shedden K, Hankenson KD, Woolf PJ. GAGE: generally applicable gene set enrichment for pathway analysis. BMC Bioinformatics 2009;10:161.
[32] Ashburner M, Ball CA, Blake JA, Botstein D, Butler H, Cherry JM, et al. Gene ontology: tool for the unification of biology. The Gene Ontology Consortium. Nat Genet 2000;25:25–9.
[33] Gene Ontology Consortium. The Gene Ontology resource: enriching a GOld mine. Nucleic Acids Res 2021;49:D325–34.
[34] Kanehisa M. Toward understanding the origin and evolution of cellular organisms. Protein Sci 2019;28:1947–51.
[35] Kanehisa M, Goto S. KEGG: kyoto encyclopedia of genes and genomes. Nucleic Acids Res 2000;28:27–30.
[36] Kanehisa M, Furumichi M, Sato Y, Ishiguro-Watanabe M, Tanabe M. KEGG: integrating viruses and cellular organisms. Nucleic Acids Res 2021;49:D545–51.
